# Using a multiscale lidar approach to determine variation in canopy structure from African forest elephant trails

**DOI:** 10.1101/2023.08.25.554381

**Authors:** Jenna M. Keany, Patrick Burns, Andrew J. Abraham, Patrick Jantz, Loic Makaga, Sassan Saatchi, Fiona Maisels, Katharine Abernethy, Christopher Doughty

## Abstract

Recently classified as a unique species by the IUCN, African forest elephants (*Loxodonta cyclotis*) are critically endangered due to severe poaching. With limited knowledge about their ecological role due to the dense tropical forests they inhabit in central Africa, it is unclear how the Afrotropics would change if forest elephants were to go extinct. Although their role as seed dispersers is well known, they may also drive large-scale processes that determine forest structure, through the creation of elephant trails and browsing the understory and allowing larger, carbon-dense trees to succeed. Multiple scales of lidar were collected by NASA in Lopé National Park, Gabon from 2015-2022. Utilizing two airborne lidar datasets and one spaceborne lidar in an African forest elephant stronghold, detailed canopy structural information was used in conjunction with elephant trail data to determine how forest structure varies on and off trails. Forest above elephant trails displayed different structural characteristics than forested areas off trails, with lower canopy height, canopy cover, and different vertical distribution of plant mass. Less plant area density was found on trails at 1 m in height, while more vegetation was found at 12 m, compared to off trail locations. Trails with previous logging history had lower plant area in the top of the canopy.

## INTRODUCTION

African forest elephants (*Loxodonta cyclotis*) are critically endangered (Gobush et al., 2021), facing severe threats from poaching and deforestation with estimated population losses of 62% in central Africa from 2002-2011 (Maisels et al., 2013). Despite their recent, broad-scale decline and listing as a separate species in 2021 (Hart et al., 2021), forest elephants are understudied due to poor visibility in African tropical forests. Specifically, the forest elephant’s ecological role is severely understudied in comparison to the African savanna elephants (*Loxodonta africana*), whose role as a keystone species has been understood for decades (Bond, 1994). Previous work identified forest elephants as important seed and lateral nutrient dispersers through plant digestion, defecation, urination and the ingestion of mineral rich water and soil (Doughty et al., 2016; Metsio Sienne et al., 2014; Blake et al., 2009). However, their influence on forest structure is less clear.

From select studies, it is hypothesized that forest elephants impact forest structure through their feeding preferences. As generalist browsers, forest elephants modify their environment by trampling, ingesting, and shaking vegetation to access preferred fruits (Maisels et al., 2002). It is proposed that through browsing in the understory, small saplings are consumed which allows large, woody trees to succeed (Terborgh et al., 2016). For example, Berzaghi et al. (2023) collated >100,000 records of elephant feeding preferences in central Africa and discovered that low wood density vegetation was consumed at significantly higher rates than high wood density tree species. On the other hand, fruits from larger trees were preferred by forest elephants. By feeding on low wood density plants but fruit from larger trees, elephants promote forests with higher biomass. It is estimated that if tree species preferentially dispersed by elephants were replaced by other species, above ground carbon could decrease by up to 12% in central Africa (Berzaghi et al., 2023).

One of the most obvious ways forest elephants impact their environment is by the creation of trails through the forest (Vanleeuwe and Gautier-Hion 1998; Blake and Inkamba-Nkulu 2004). These paths are usually 0.5 to 0.9 m wide on average (Blake and Inkamba-Nkulu 2004; Vanleeuwe and Gautier-Hion 1998), allowing elephants to access important resources such as streams, mineral-rich areas (bais), and high-priority fruit trees (Blake and Inkamba-Nkulu 2004; Benitez and Queenborough 2021). Through consistent use, structural changes in the understory have been found along elephant trails in the Republic of the Congo (Blake and Inkamba-Nkulu 2004; Valeeuwe and Gautier-Hion 1998), the DRC (Inogwabini et al. 2013), and Uganda (Benitez and Queenborough 2021). Vanleeuwe and Gautier-Hion (1998) found that forest elephants created ‘boulevard trails’ for migration which cut through all forest types and ‘foraging trails’ which wound through the thick Marantaceae and Zingiberaceae (terrestrial herbaceous vegetation). Elephant trail networks are dense (9.82 trails/km in Ndoki National Park; Blake and Inkamba-Nkulu, 2004) and probably last for hundreds of years as elephants pass on their mental maps of the forest to their offspring (Haynes, 2006).

Additional fauna—including humans, maintain elephant trail systems and utilize them to access rivers, bais, and hard-to-access regions of the forest (Blake, 2002; Remis and Robinson, 2020). Indigenous tribes such as the BaAka, have used elephant trails for hunting and travel for centuries (Remis and Robinson, 2020). Elephant trails in Gabon were found to act as natural firebreaks along forest-savanna boundaries, assisting in the protection of forest interiors (Cardoso et al., 2020). Understanding the complexities of vegetation structure surrounding these trails could shed light on forest elephant feeding behaviors, dispersal patterns, space use, and role as ecosystem engineers—all of which are still relatively unknown.

Lidar (light detection and ranging) sensors have been used to measure forest structural properties, such as cover and canopy height (Dubayah and Drake, 2000). Energy pulses returned from the ground and vertical sub-canopy vegetation structure are digitized and converted into a 3D representation of topography and plant biomass distribution. Lidar has proven effective in modeling habitat preferences, understanding predator-prey dynamics, and determining species richness in relation to both vertical and horizontal structure (Goetz et al., 2010; Davies and Asner, 2014; McLean et al., 2016). To our knowledge, lidar has never been used to determine forest elephants’ impact on the structure of forests in the Afrotropics.

Here, we use remotely sensed lidar to investigate vegetation structure at broad spatial scales along forest elephant trails in Lopé National Park, Gabon. This study aims to characterize the role forest elephants play as ecosystem engineers through the creation of trails, specifically by using lidar to quantify their impact on canopy structure. Multiple lidar collections have taken place over the last 8 years in Lopé National Park, Gabon, a protected area with one of the highest elephant densities in central Africa (0.93 elephants/km^2^; Bezangoye and Maisels, 2010). Utilizing elephant trail geolocation data and three scales of lidar (2x aircraft, 1x satellite) from NASA, questions regarding how elephants affect canopy structure along trails can be addressed. Specifically, our research questions are:

1. Can changes in forest structure associated with elephant trails be detected with lidar? If so, which lidar sensors can detect structural variation?
2. How do lidar-assessed canopy structure metrics (e.g., canopy height, canopy cover, vegetation area index, height of median energy) vary with distance from elephant trails?

## METHODS

### Study Area

Lopé National Park covers 4960 km^2^ near the center of Gabon (0° 10’S 11° 35’ E; Figure 1) and was established as a nature reserve in 1946 and as a UNESCO World Heritage Site in 2007 for its natural and archeological richness. The park is dominated by closed canopy tropical forest with a forest-savanna mosaic located in the northern section. Forest types include mature, Marantaceae, gallery, and young forest (due to savanna colonization). The most dominant vegetation type in Lopé is mature forest (63.7%), which comprises medium to large trees with very dense canopy cover and a limited understory (White and Abernethy, 1997). Marantaceae forest is the second most prevalent forest type characterized by dense understory dominated by Marantaceae (arrowroot family) and other herbaceous vegetation preferred by elephants and gorillas, with large trees dominating the canopy (White and Abernethy, 1997). Gallery forests are located along water courses; small woodland patches (“bosquets”) are located in the savannas. Both typically have shorter canopies and sparser understories than mature or Marantaceae forests. Lopé Faunal Reserve was selectively logged from the 1960’s to the early 2000’s for Okoumé (*Aucoumea klaineana*) trees and over 50 other hardwood species with removal rates <2 trees/ha, prior to becoming a national park (White and Taylor, 1994). Since the gazettement of the area as a National Park in 2002, there has been limited human influence in the park due to a low density of human settlement in this region of Gabon. Eight settlements occur along the northern and north-eastern park boundaries (<4,000 people in total) and some indigenous communities use the interior of the southern region of the park (Rakotonarivo et al., 2021).

**Figure 1.**
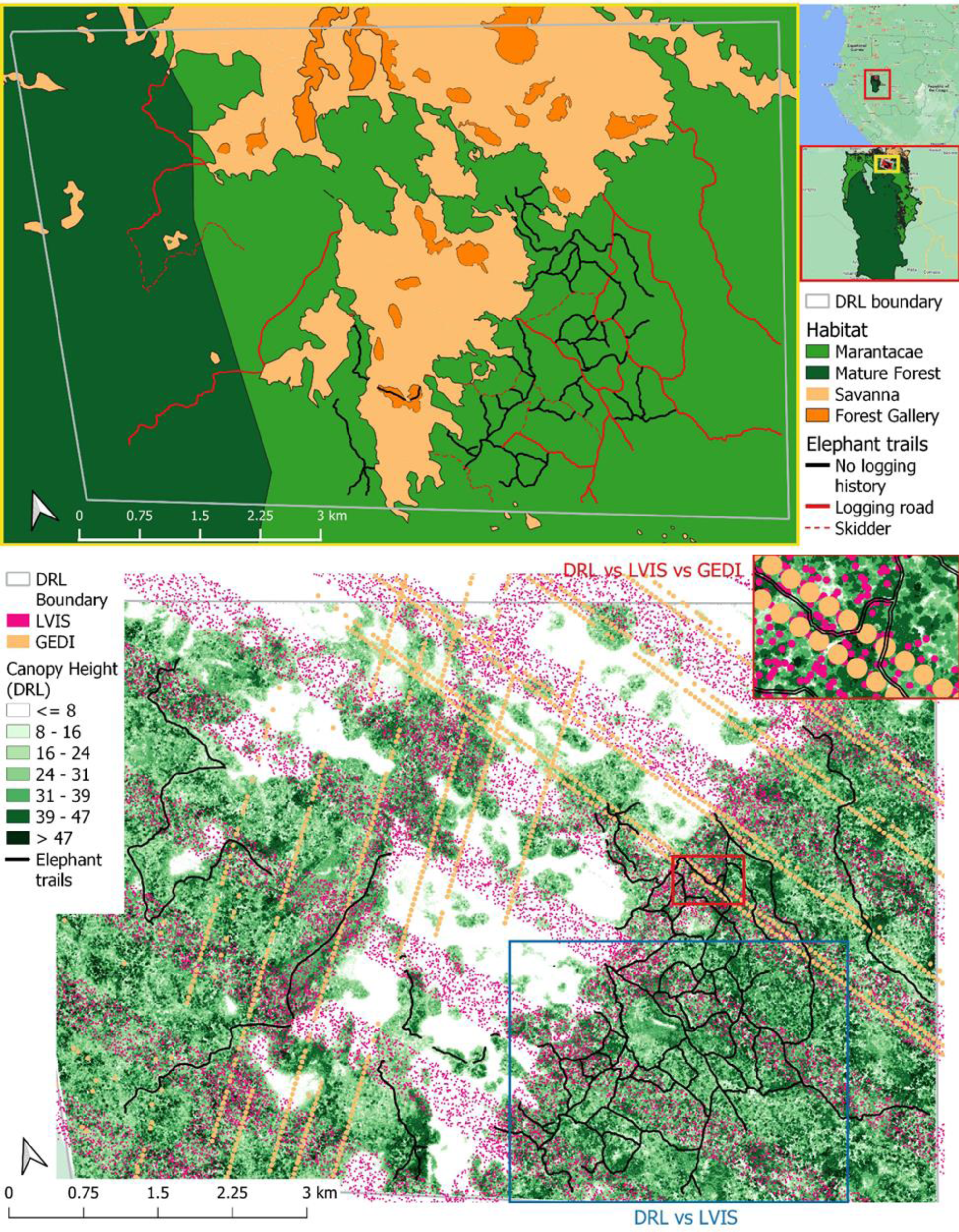
Study area within Lopé National Park, Gabon. Habitat types and elephant trail history are indicated in the top panel, while the three lidar datasets are shown in the bottom panel with overlapping elephant trails (black). The blue inset map highlights the region where the two airborne lidar datasets have the most overlap (DRL vs LVIS) and the red inset highlights where all three lidar datasets (DRL vs LVIS vs GEDI) overlap the most elephant trails.

### Elephant Trail Data

Fifteen km of elephant trails were surveyed and geolocated in Lopé National Park in January 2022. Trail data were collected with a handheld GPS (Garmin 66i), which has a standard 3-meter accuracy. Tracklogs were then downloaded onto Garmin BaseCamp and converted to shapefiles in QGIS for analysis. Width of trails at breast height (∼1.4m) and on the ground were measured and recorded every 200 m along trails. Trail edges on the ground were determined by the presence of bare ground, while trail edges at breast height were recorded where the vegetation subsided on either side. These data were supplemented by an existing dataset with 62 km of georeferenced elephant trails from Gabon’s National Park Agency (ANPN), collected between 1986 and 2010 and regularly refined as path locations changed and GPS technology advanced (White, 1995; SEGC unpublished).

Elephant trail history was verified by K. Abernethy (Associate Researcher for the National Centre for Research in Science and Technology in Gabon) and L. Makaga (Lopé Research Station Manager) as either: (a) elephant-engineered (b) previous logging road (c) previous skidder trail. All trails are now elephant trails, while some were previously created or used by the logging industry to access the forest more easily. Skidder trails were used to extract logs, with some small trees removed but not clear-cut or bulldozed. Previous logging road trails were clear-cut and bulldozed to achieve an understory opening of >5 m, allowing heavy equipment to enter the forest. Elephant-engineered trails were never widened or logged but have been maintained by elephants. To simplify these classifications, trails that were once logging roads were deemed ‘elephant-and-human engineered’, while trails maintained by elephants only are ‘elephant-engineered’. Skidder trails were removed from the analysis due to a small sample size. Although all logging in the study area of the park ceased 40-50 years ago, it is vital to take the effect of logging into account due to its known legacy in affecting forest structure (Hall et al., 2003).

### Lidar Data

We used lidar data acquired from three different sensors, with spatial resolution ranging from ∼ 1 to 25 m (Table 1). The first dataset was acquired in July 2015 as a discrete return point cloud using a low-flying rotary platform, providing 54 km^2^ of wall-to-wall coverage (Discrete Return Lidar or DRL; Silva et al. 2018). The second dataset was acquired in March 2016 as waveform lidar using NASA’s Land, Vegetation, and Ice Sensor (LVIS) mounted on a King Air B-200 flown at 7.3 km (LVIS; Blair et al., 1999). The third dataset was acquired between April 2019 and October 2022 as waveform lidar using the Global Ecosystem Dynamics Investigation sensor mounted on the international space station (GEDI; Dubayah et al., 2020). Both LVIS and GEDI data return individual ‘shots’ instead of wall-to-wall coverage, ranging from 20 (LVIS) to 25 m (GEDI) in diameter. Figure 1 shows the location of the three lidar datasets in relation to known elephant trails surveyed in Lopé National Park. Table 1 describes the spatial resolution, instrument specifications, and data collection timeline for each lidar acquisition. With the highest spatial resolution of all lidar in the study at 1 m, the DRL dataset provides wall-to-wall coverage of the study area (Figure 1). The DRL and LVIS data were validated with field plots and compared in 2018 for structural metrics (Silva et al., 2018). LVIS canopy cover and vertical profile products (plant area volume density or PAVD) were downloaded from Oak Ridge National Laboratory Distributed Active Archive Center (ORNL DAAC; Tang et al., 2018). GEDI products were downloaded from the Land Processes DAAC. Canopy height (RH98), canopy cover (CC), and plant area (PAVD) were extracted from GEDI’s L2A and L2B data from April 17, 2019 to October 25, 2022. All GEDI shots were then collocated using the DRL to ensure the highest accuracy in geolocation.

**Table 1.** Lidar data specifications from both airborne and spaceborne sensors.

| *Name* | *Type* | *Platform + Instrument* | *Footprint Width* | *Vertical / Horizontal Geolocation Accuracy* | *Pulse Density* | *Acquisition Date(s)* |
| --- | --- | --- | --- | --- | --- | --- |
| DRL | Airborne Discrete Return Point Cloud | EC 135 Helicopter + Riegl VQ480U (1550 nm) | < 0.1 m | < 1 m / < 1 m | 78,000 pulses/ha | July 2015 |
| LVIS | Airborne Full Waveform | NASA Langley King Air B-200 Airplane + LVIS laser (1064 nm) | 20 m | ~ 1 m / 0.1 m | 4.6 points/ha | March 2016 |
| GEDI | Spaceborne Full Waveform | ISS + 3 HOMER lasers (1064 nm) | 25 m | ~ 10 m / ~ 1 m | 0.23 points/ha | April 2019 to Oct. 2022 |

### Data Analysis

#### A) Elephant trail data

Elephant-engineered and elephant-and-human engineered trails in Marantaceae forests were used in the lidar analysis. These were selected as Marantaceae is the dominant habitat type in our study area, and skidder trails had a small sample size (mature forest and forest gallery results are found in the supplementary material). All elephant trails were clipped to exclude savannas. A 2 m diameter buffer (1 m from trail center) was used for ‘on trail’ analyses to account for handheld GPS positional uncertainty. While GPS uncertainty fluctuates, elephant trails are on average 0.5-0.9 m in width. Therefore, in order to detect any understory changes in vegetation, a 2 m buffer was chosen. Additional 10 m buffers were created from 10 to 60 m from trails and used to compare on and off trail vegetation. Waterways were removed from all trail shapefiles using a 50 m diameter (25 m from center) stream buffer (Figure 2).

**Figure 2.**
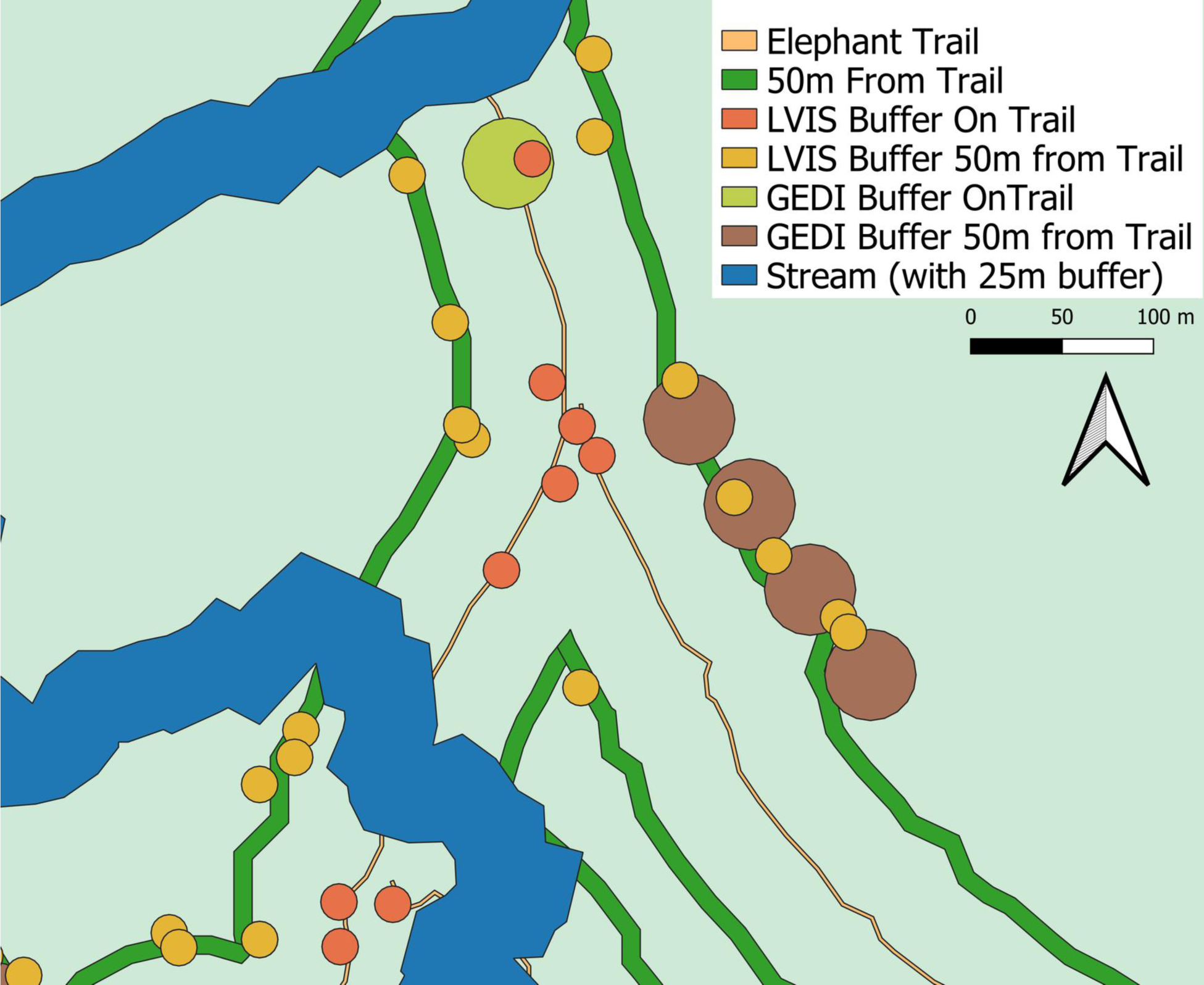
Visualization of buffered trail and lidar data used in the structural analyses.

#### B) Lidar Processing

The discrete return lidar were clipped to both on and off trail buffers (segment tool) and filtered using the statistical outlier filter with 2 standard deviations in CloudCompare (v.2.13alpha). Any additional erroneous points located above top of canopy height or below the ground were manually removed with the segment tool. Within R (version 4.2.1), the lidR, sp, and lidRmetrics packages were used extensively to process and manipulate the point cloud (Roussel et al., 2020; Roussel and Auty, 2023). A digital terrain model (DTM) was created from the cleaned point cloud using the kriging method (k=10L), which was then used to normalize the height of all lidar returns. An additional manual filter of the normalized point cloud was performed to ensure a clean point cloud without erroneous points (0 < Z< 65). LVIS and GEDI point data were buffered by their respective footprint widths (20 m in diameter for LVIS and 25 m for GEDI; Figure 2). These buffers were then used to clip the wall-to-wall DRL pointcloud for the multiscale lidar comparisons (DRL vs. LVIS and DRL vs. LVIS vs. GEDI) to ensure direct overlap between all lidar scales for forest structure comparisons. GEDI and LVIS footprints that had at least 50% overlap with elephant trail buffers were selected for subsequent canopy structural analyses.

#### C) ‘On Trail’ and ‘Off Trail’ Forest Structural Comparisons

We used a variety of canopy metrics to compare vegetation structure on and off elephant trails (Table 2). First, leaf area density (LAD) or plant area volume density (PAVD; Dubayah et al., 2021) were used to understand how the vertical structure profile differed on and off trails. LAD and PAVD quantify the distribution of both woody and foliar material across discrete height bins (Table 2). LVIS PAVD is discretized into 1 m height bins, while GEDI PAVD is discretized into 5 m height bins. LAD was calculated in the subsequent DRL using the lidR package (Roussel et al., 2020) from the cleaned and normalized point cloud into 1 m height bins using a standard extinction coefficient of foliage of 0.5 and a clumping index of 1 (also used for LVIS and GEDI). Although LAD and PAVD are calculated using different equations, they are both indicators of the vertical distribution of plant area in the forest.

**Table 2.** Canopy structural metrics descriptions and references.

| Metric | Description | Reference |
| --- | --- | --- |
| Leaf Area Density (LAD/PAVD) | Leaf area per unit of volume | Bouvier et al. 2015 |
| Max Canopy Height (CH) | Maximum of canopy height | Drake et al. 2001 |
| Height of Median Energy (HOME) | Median of returned energy | Drake et al. 2001 |
| Vertical Distribution Ratio (VDR) | $VDR = [CH - HOME] / CH$<br>An index ranging from 0 to 1 depicting the vertical distribution of plant matter | Goetz et al. 2007 |
| Total Vegetation Area Index (VAI) | Sum of LAD for all height bins | Bouvier et al. 2015 |
| Canopy Cover Fraction (CCF) | 1- Deep Gap Fraction | LaRue et al. 2020 |

The maximum canopy height is equated with the LVIS and GEDI RH98 metric - the height (relative to the ground) at which 98% returned energy is reached (Table 2). RH98 represents the top of the canopy and is not as sensitive to atmospheric noise such as fog or very low clouds. Canopy height is an indicator of early forest successional stage and above ground biomass (Drake et al. 2001). RH50 is also referred to as the height of median energy (HOME), which indicates where the bulk of vegetation is located vertically. The vertical distribution ratio (VDR; Goetz et al. 2007) equals (RH98 – RH50) / RH98. VDR is normalized by RH98 to indicate the relative distribution of biomass within the vertical profile. High VDR values (closer to 1) are associated with bottom-heavy vertical profiles (mid successional) while lower values (closer to 0) are associated with top heavy vertical profiles (young and old forests). Finally, canopy cover fraction (CCF) was calculated to determine the amount of forest cover within each buffer. Canopy cover fraction is ecologically important as it determines light availability, water interception, and temperature regulation.

Violin plots for all canopy metrics were created and Wilcoxon tests were used to compare each metric for on and off trail forests. Finally, LAD vertical bins were tested for normality using Kruskal Wallace tests. As the data were not normally distributed, Wilcoxon tests were used to determine differences in LAD/PAVD for each vertical height bin (either 1 m for DRL and LVIS or 5 m for GEDI).

## RESULTS

### Canopy Structural Metrics

Elephant-engineered trails and elephant-and-human trails had an average ground width of 88.8 cm and 123.7 cm respectively (Supp. Table 1). Average width at breast height increased for both trail types, with elephant-engineered at 218.9 cm and elephant and human at 287.1 cm (Supp. Table 1). When averaging leaf area density from 1-4 m in height with the DRL, less LAD is found on trails than off (Figure 3). However, this increase is relatively small with LAD values generally around 0.6 m^2^m^-3^. Furthermore, both elephant-engineered and elephant-and-human trails had significantly less LAD on trails than off at the 1 m vertical height bin (Figure 3).

**Figure 3.**
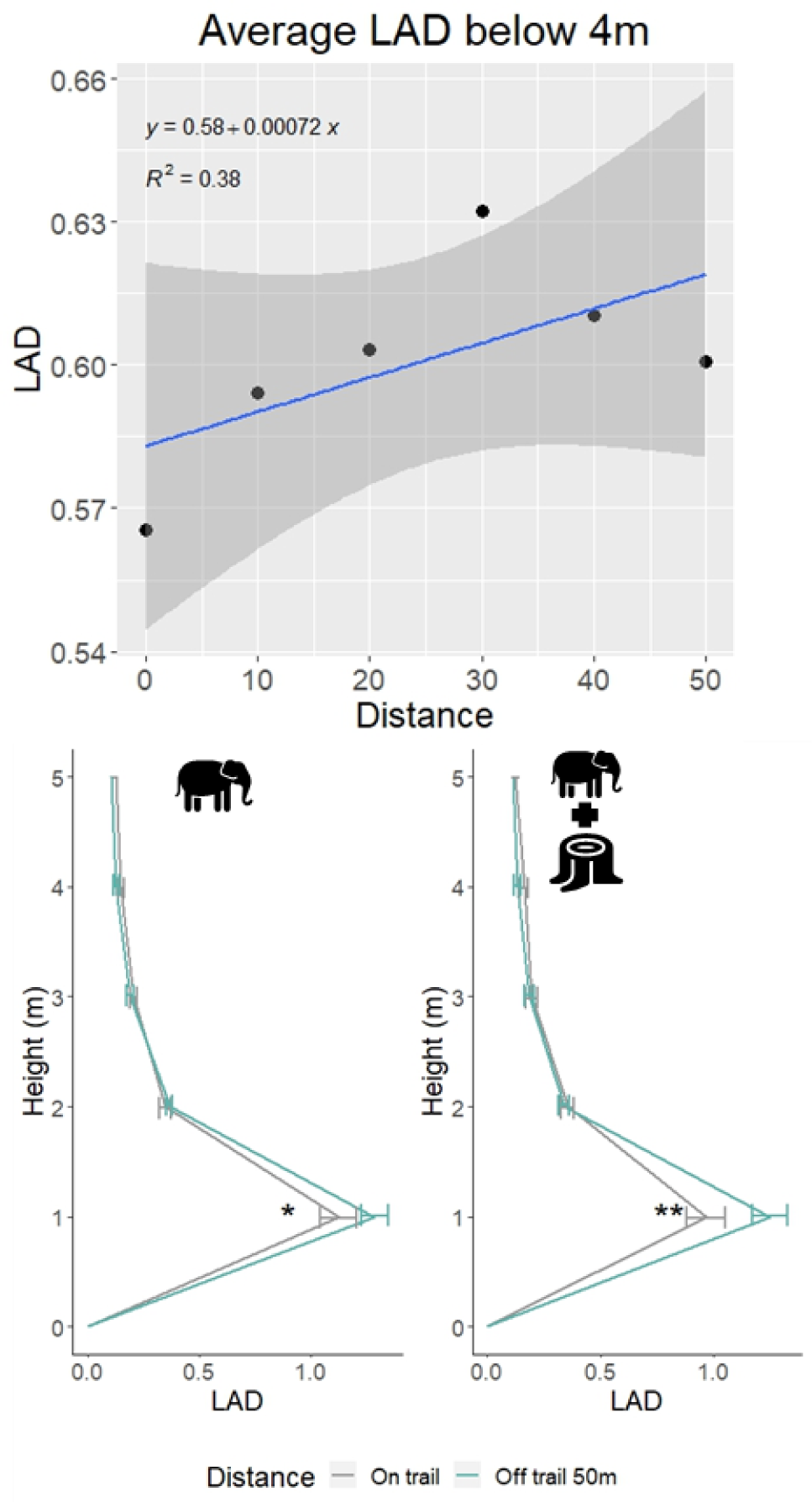
DRL average leaf area density from 1-4m on elephant engineered trails in Marantaceae forests fitted with a linear trend line with standard error. The trend line accounts for 38% of the variation. Left: elephant engineered only; right: elephant and human engineered trails.

All ‘on trail’ and ‘off trail’ buffers were analyzed for forest structure using the DRL, as it provided full coverage of the study area. From these analyses, max canopy height was significantly higher 50 m from trails than on both elephant-engineered (p <0.001) and elephant-and-human trails (p<0.01) (Figure 4). No difference was found between HOME or VDR on elephant-engineered trails, however elephant-and-human had higher HOME values off trail (p<0.01) and lower VDR values off trail than on (p<0.05). Total VAI was higher off trails than on elephant-and-human trails (p<0.01), with no difference observed for elephant-engineered trails.

**Figure 4.**
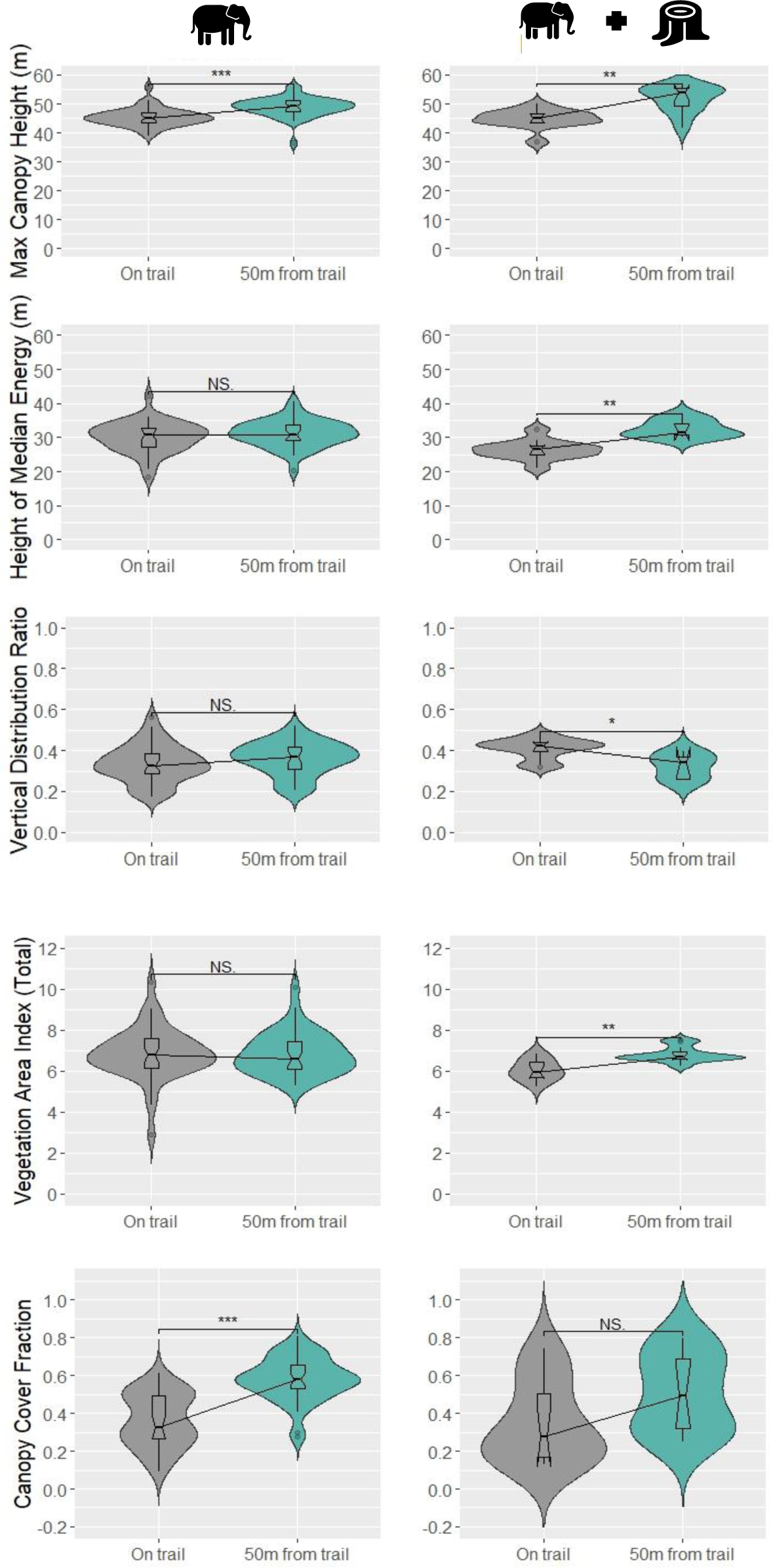
Total VAI, CCF, Max CH, HOME, and VDR violin plots from DRL on elephant trails and 50m from elephant trails. Comparisons using elephant engineered trails are indicated on the left panels, while elephant and human engineered trails are indicated in the right panels. Notched box plots are within each violin plot with trend lines connecting median values between on and off trail. Statistical comparisons using the Wilcoxon test between on and off trail are indicated as not significant (NS), one asterisk (p<0.05), two asterisks (p<0.01), or three asterisks (p<0.001).

Elephant-engineered trails showed significantly higher canopy cover 50 m from trails than on them (p<0.001), and no significant difference was found in cover between on and off elephant- and-human trails. When comparing the vertical profiles (average leaf area density) of for on and off trail forest, elephant-engineered trails had higher LAD at the 1, 12, 47, 48, and 49 m height bins (Figure 5). Elephant-and-human trails had less LAD at 1 m and 30-49 m heights and more LAD at 16, 17, and 18 m heights than off trail forest.

**Figure 5.**
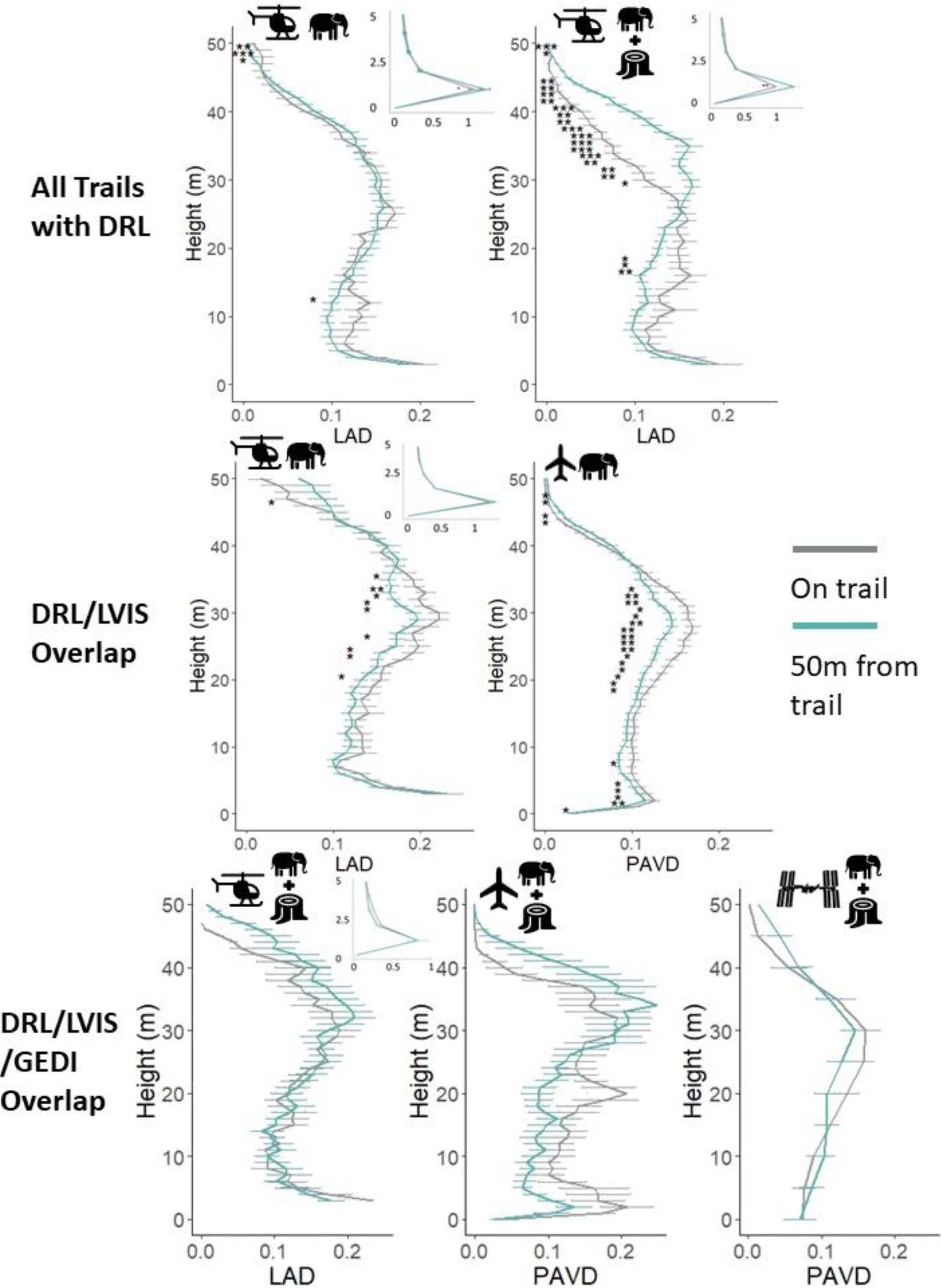
(Top) Average LAD profiles from DRL for elephant-engineered trails and elephant and human engineered trails with standard error bars. (Middle) Average LAD and PAVD profiles with standard error bars from overlapping DRL and LVIS shots on elephant engineered trails (N=219 on trail, N=355 off trail). (Bottom) Average LAD and PAVD profiles with standard error bars from overlapping DRL, LVIS, and GEDI shots on elephant and human engineered trails (N=9 on trail, N=13 off trail). Statistically significant differences from Wilcoxon tests for each height bin (1 m for DRL and LVIS and 5 m for GEDI) are indicated by one (p<0.05), two (p<0.01), or 3 (p<0.001) asterisks. Inset figures display DRL LAD values from 0-5 m.

### Comparing Multiple Lidar Scales

To determine what scale of lidar can be used to detect changes in forest structure on elephant trails, overlapping lidar data were compared on and off trails. Following the wall-to-wall DRL, airborne LVIS data had the most overlap with the study area (blue box in Figure 1). Therefore, the DRL and LVIS were used to compare canopy structure on and off elephant-engineered trails (Figure 5). Significant differences were observed in plant area density from DRL height bins at 22-36 and 48 m (Figure 5). Almost all height bins were significantly different between on and off elephant-engineered trails from LVIS, except from 10-20 and 35-40 m (Figure 5). A similar pattern is seen in the DRL vs LVIS comparison for the elephant-and-human engineered trails (Supp. Figure 1). The lidar scale with the least coverage was GEDI, with 9 shots on elephant- and-human trails and 13 off. There were no GEDI shots on elephant-engineered trails. Using all three lidar scales, no significant differences were found between on and off trail forest structure (Figure 5).

Due to smaller lidar overlap in mature forests and forest galleries, limited canopy structural analyses were completed for those habitat types. Using the two airborne lidar datasets (DRL and LVIS), elephant-and-human trails in mature forests depicted similar canopy structure to those in Marantaceae forests with higher leaf area density on trails in from 1-15 m and lower leaf area on trails from 30-45m (Supp. Figure 2). Elephant-engineered trails in forest galleries also showed similar trends to those in Marantaceae forests (Supp. Figure 3). GEDI shots were either absent or limited (N<5) for mature forest and forest galleries.

## DISCUSSION

### African Forest Elephant Trails Impact Canopy Structure

Elephant trails in the Marantaceae forests of Lopé National Park, Gabon appear to impact forest structure by reducing canopy cover and canopy height, while changing the vertical leaf area density structure. Less leaf area is found 1 m above the ground for both trail types, while more leaf area is found at 12 m above the ground for elephant-engineered trails, and at 16, 17, and 18 m in height for elephant-and-human trails. Leaf area is significantly less on elephant-and-human engineered trails from 30-49 m in height. With similar HOME and VDR metrics, the vertical distribution of plant area is similar both on and off of elephant-engineered trails, however canopy height and cover are smaller on trails (Figure 4). Through the creation and maintenance of trails, it appears that African forest elephants impact canopy structure by creating less canopy cover and lowering canopy height of vegetation while keeping total VAI the same. Total VAI was similar both on and off elephant-engineered trails indicating that the higher plant area in the lower vertical bins made up for a shorter canopy.

When comparing forest structural changes from elephant-engineered trails to trails altered by the logging industry (through clear-cutting and bulldozing), considerably more impact on the canopy is observed. The legacy of selective logging is noticeable in our study, showing that after ∼50 years since logging subsided, the overall vertical structure remains affected. The widening and compacting of trails during tree harvesting produced long lasting effects by lowering canopy height, total VAI, HOME, and canopy cover (Figure 4). These structural changes are in line with previous research, showing that logged forests in central Africa have altered ecosystem composition from the loss of larger trees and increased diversity in the understory from higher light penetration (Hall et al., 2003; Sullivan et al., 2022). Yet, the intertwining influences of both elephant and logging disturbance to vegetation on trails are novel. While the effects of logging in the Afrotropics are well documented (Cazzolla Gatti et al., 2015), our study suggests that African forest elephants create a lighter disturbance in the canopy—acting as a ‘logging light’ ecosystem engineer. To understand these effects in more depth, we identified two hypotheses on how elephant trail vegetation structure is established.

To address our findings of higher plant area from 5-30 m above the ground, we hypothesize that elephants create trails in the thickest Marantaceae to gain access to better browse, while simultaneously promoting understory growth through increased canopy gaps. While some trails could be direct paths moving toward a water source or fruiting tree, most others could be created to access the Marantaceae roots—a preferred browse by forest elephants (White et al., 1993). In a similar manner to logging, additional gaps in the forest caused by forest elephant browsing and soil compaction would allow more light to penetrate the canopy and reach the ground (Struhsaker et al., 1996; Kamga et al., 2022), which increases growth of understory vegetation. Added light to the lower and mid canopy promotes the expansion of light seeking fast-growing saplings. Consequently, a feedback effect consisting of elephants consuming vegetation along trails while also promoting more growth is possible. Mount Cameroon National Park has a natural elephant exclosure area (from lava flows), allowing researchers to test forest structure in elephant and non-elephant areas of the park. In this unique environment, forest elephants promoted canopy gaps and fostered a more heterogeneous forest, which in turn increased the diversity of insects and birds (Maicher et al., 2020; Kamga et al., 2022). Our findings support this hypothesis by showing that both elephant-and-human and elephant only trails have lower canopy cover (Figure 4) and higher plant area in the understory (Figure 5).

Our second hypothesis is that increased soil fertility along trails from animal use encourages more fast-growing vegetation to thrive along elephant trails. The addition of seed deposition and disturbance from forest elephants might promote more fertile growing conditions near elephant paths. Megaherbivores are known to disproportionately affect nutrient availability and forest fertility through the deposition of feces and urine (Wolf et al., 2013; Doughty et al., 2016; Stanbrook et al., 2018; Enquist et al., 2020). Little is known about forest elephant’s use of space in regard to departing from trails to enter the forest interior, however Inogwabini et al. (2013) found less elephant dung with increasing distance from trails, suggesting they typically stay along the established paths, although the sample size was small. Increased nutrient deposition along elephant trails could therefore promote increased fast-growing vegetation from higher soil fertility. Finally, the effects of higher soil nutrients could contribute to lower canopy height on trails, which is in line with previous findings associating lower aboveground biomass with higher N and P concentrations in the tropics (Unger et al., 2012).

### Determining What Scale of Lidar is Needed to Detect Elephant Trails

It is apparent that lidar can distinguish changes in forest structure from elephant paths, but what lidar is necessary to determine these differences? As expected, our multiscale lidar analysis found the highest resolution airborne lidar (DRL) was the best scale to determine fine-scale changes in structure. Furthermore, airborne full waveform lidar detected variation in plant area better than spaceborne lidar. Through NASA’s continued investment in collecting LVIS in Gabon, additional data will be available in the future. With GEDI’s 25 m footprints and current sampling density, analyzing animal trails is difficult. However, with more data collections on future missions, near global lidar could prove important for understanding how animals such as forest elephants impact structure.

### Limitations and Uncertainties

Additional limitations to this study include anthropogenic influences, collection time differences, and geolocation uncertainties for the trail and lidar data. With average widths of 88 cm on elephant-engineered trails, a 3-meter uncertainty buffer on the GPS units could produce changes in the lidar results, making it difficult to find the understory vegetation gap along the path. Each lidar dataset has accompanying GPS errors which could impact our results as well. Silva et al. (2018) compared the DRL and LVIS collected in Lopé and found some large differences in Z values on a footprint scale, mainly due to complex topography. When compared across larger scales, variation decreased and no significant differences were found. With the 3 lidar comparison, collocating the GEDI shots with the DRL instilled greater confidence in our findings as it accounted for known geolocation issues with GEDI. Nonetheless, using GEDI to determine fine-scale changes in vegetation from animal paths is difficult. UAV lidar could be highly useful when studying small trails in thick Marantaceae forests and should be explored in future research. The expansive spatial coverage of GEDI could be especially useful in larger study areas with no available airborne lidar collections.

### Conclusion

Using remotely sensed lidar, we were able to determine canopy height, canopy cover, and vertical structural differences between forest along elephant trails and the surrounding areas in Lopé National Park, Gabon. We also showed that a structural signature exists on logging roads now used as elephant trails, even 50 years beyond the abandonment of logging activity. These findings are novel, as lidar has never been used to quantify how large animals such as African forest elephants influence canopy and vegetation structure through the creation of trails (to our knowledge). Nor has the persistence of the structural impacts from selective logging been followed on such a long time scale. Although the highest resolution lidar detected more variation in forest structure than LVIS and GEDI, increased coverage of full waveform lidar in the African tropics is imperative to broaden our understanding of elephants’ ecological role. Future studies on forest elephant trails in these habitat types would benefit from measuring light availability, characterizing Marantaceae thickness, and classifying the trail use with direct measurement using camera traps. Additional research is needed to understand elephant preferences and motivations (e.g. fruiting trees, water courses, bais) to create their trail networks in a variety of habitat types. The full effect elephants have as engineers and trailblazers of the Gabonese forests is still under investigation, but it is clear they play a role in canopy height, canopy gaps, and the vertical distribution of plant mass.

## ACKNOLWEDGMENTS

We are grateful to the Gabonese Agence Nationale des Parcs Nationaux (ANPN), Centre National de la Recherche Scientifique et Technique (CENAREST), and Wildlife Conservation Society for their permission and logistical support to conduct research in Lopé National Park. We also thank the National Aeronautics and Space Administration (grant 80NSSC21K1636) for their financial support through the Future Investigators in NASA Earth and Space Science and Technology (FINESST) program. Finally, we thank Brice Momboua, Heddy Milamizokou, and Pacôme Dimbonda for their incredible kindness and assistance in the field.

## DATA ACCESSIBILITY

African forest elephant trail data is not publicly accessible as it is sensitive information of a critically endangered species. The discrete return lidar is maintained by Sassan Saatchi at the Jet Propulsion Lab and requires permission to access it. Both full waveform lidar datasets (LVIS and GEDI) are publicly available through the ORNL DAAC (https://daac.ornl.gov) and LP DAAC (https://lpdaac.usgs.gov).

## AUTHORS’ CONTRIBUTIONS

JK, AA, PJ, KA, and CD conceived of the idea and study design; SS, JK, and LM collected the data; JK and PB analyzed the data; JK, CD, FM, and KA led the writing of the manuscript. All authors contributed to the editing process.

## AUTHORS’ DECLARATION

All authors have contributed substantially to the work and approved its submission for publication to *Remote Sensing in Ecology and Conservation*. This manuscript has not been published elsewhere (in part or as a whole) nor is under consideration for publication by another journal.

