## Supplementary material for "Using a multiscale lidar approach to determine variation in canopy structure from African forest elephant trails": Supp. Table 1

Supp. Table 1. Field measurements of elephant trails in Marantaceae forests collected in Lopé National Park, Gabon, in January 2022.

| *Trail Type* | *Mean Ground Width (cm) with SD* | *Mean BH Width (cm) with SD* |
| --- | --- | --- |
| Elephant-Engineered | 88.8 ± 32.4 | 218.9 ± 60.3 |
| Elephant & Human Engineered | 123.7 ± 44.1 | 287.1 ± 86.8 |

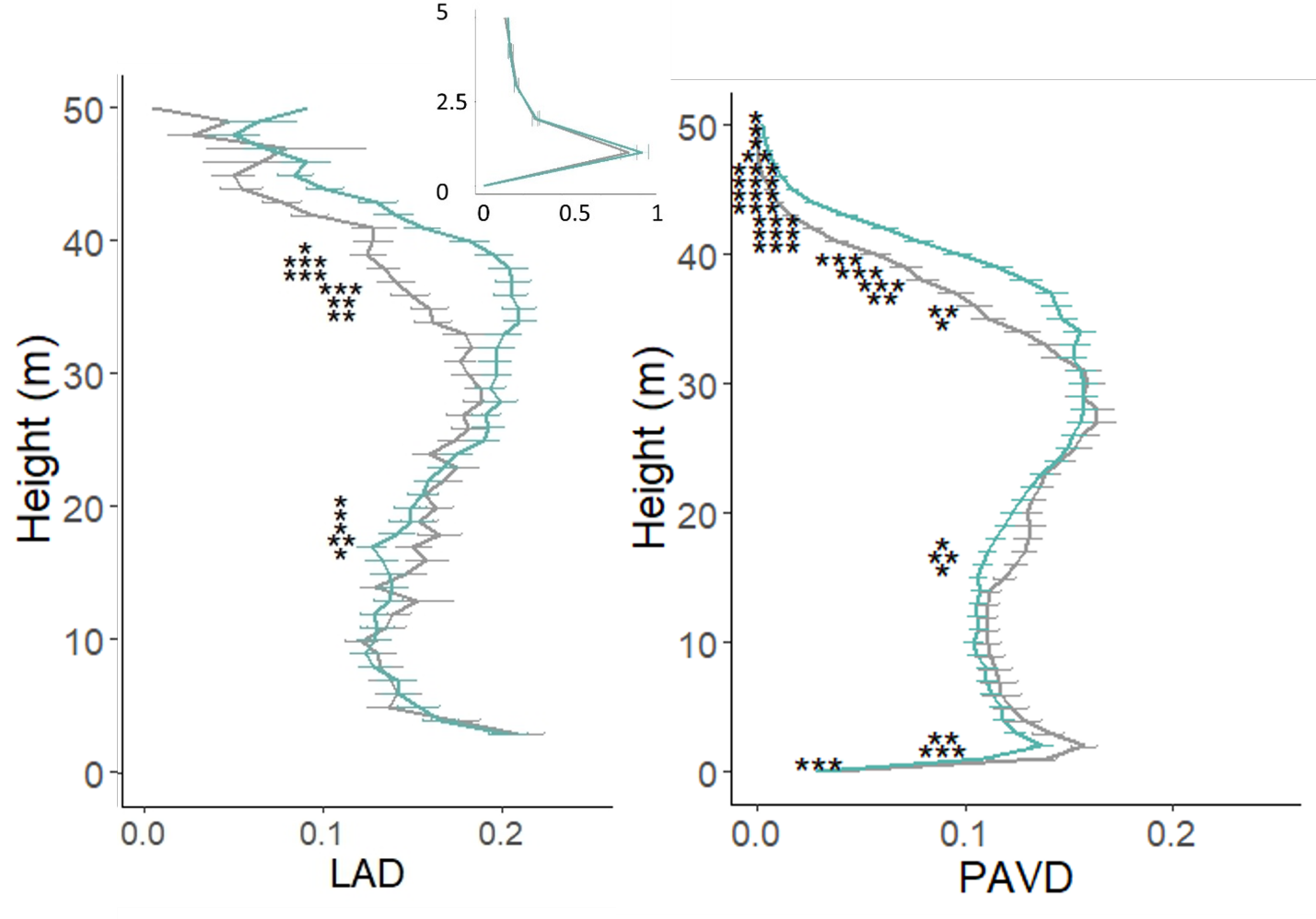

Supp. Figure 1. DRL and LVIS leaf area density comparison on elephant-and-human engineered trails in Marantaceae forests.

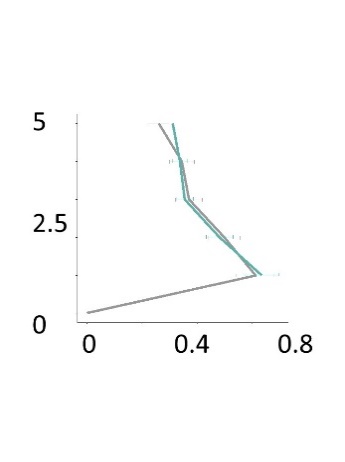

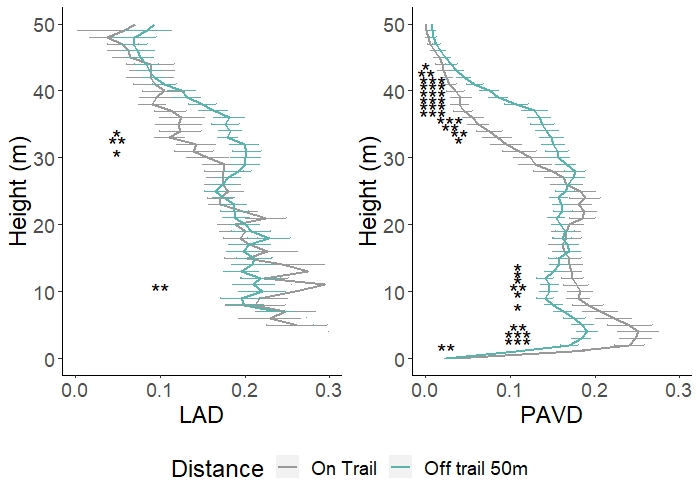

Supp. Figure 2. DRL and LVIS leaf area density comparison on elephant-and-human engineered forests in mature forests.

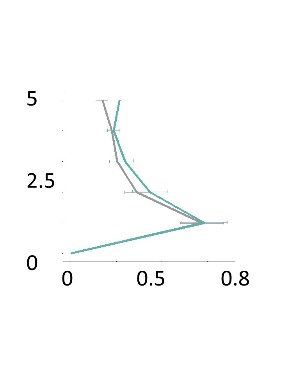

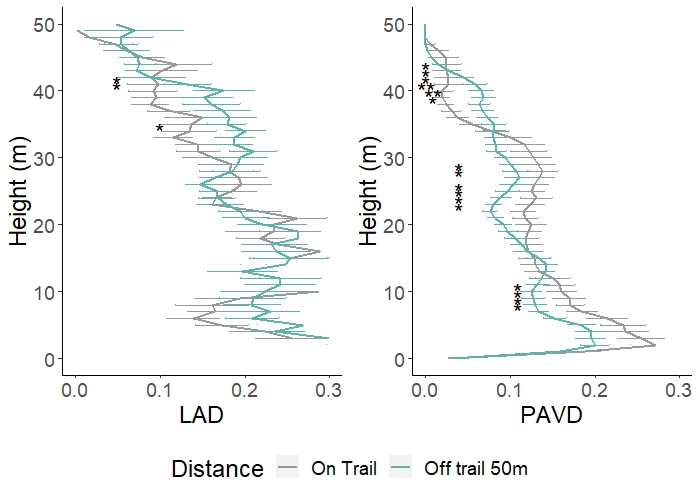
Supp. Figure 3. DRL and LVIS leaf area density comparison in forest galleries on elephant-engineered trails.
